# What makes a temperate phage an effective bacterial weapon?

**DOI:** 10.1101/2023.10.04.560906

**Authors:** M. J. N. Thomas, M. A. Brockhurst, K. Z. Coyte

## Abstract

Temperate bacteriophages (phages) are common features of bacterial genomes and can act as self-amplifying biological weapons, killing susceptible competitors and thus increasing the fitness of their bacterial hosts (lysogens). Despite their prevalence, however, the key characteristics of an effective temperate phage weapon remain unclear. Here we use systematic mathematical analyses coupled with experimental tests to understand what makes an effective temperate phage weapon. We find that effectiveness is controlled by phage life history traits – in particular, the probability of lysis, and induction rate – but that the optimal combination of traits varies with the initial frequency of a lysogen within a population. As a consequence, certain phage weapons can be detrimental when their hosts are rare, yet beneficial when their hosts are common, while subtle changes in individual life history traits can completely reverse the impact of an individual phage weapon on lysogen fitness. We confirm key predictions of our model experimentally, using temperate phages isolated from the clinically relevant Liverpool Epidemic Strain of *Pseudomonas aeruginosa*. Through these experiments, we further demonstrate that nutrient availability can also play a critical role in driving frequency-dependent patterns in phage-mediated competition. Together, these findings highlight the complex and context-dependent nature of temperate phage weapons, and highlight the importance of both ecological and evolutionary processes in shaping microbial community dynamics more broadly.

**Importance:** Temperate bacteriophage – viruses that integrate within bacterial DNA – are incredibly common within bacterial genomes. These phages are thought to act as powerful self-amplifying weapons, allowing their bacterial hosts to kill nearby competitors and thus gain a fitness advantage within a given niche. But what makes an effective phage weapon? Here we first use a simple mathematical model to explore the factors determining phage weapon utility. Our models suggest that phage weapons are nuanced and context-dependent: an individual phage may be beneficial or costly depending upon tiny changes to how it behaves, or to the bacterial community in which it resides. We then confirm these mathematical predictions experimentally, using phage isolated from Cystic Fibrosis patients. But, in doing so, we also find that another factor – nutrient availability – plays a key role in shaping phage-mediated competition. Together our results provide new insights into how temperate phage modulate bacterial communities.

## Introduction

Temperate bacteriophages (phages) provide their bacterial hosts (lysogens) with resistance to genetically similar phages through superinfection exclusion (Bondy-Denomy et al., 2016, Bordet, 1925, Lwoff, 1953), enabling their use as self-amplifying biological weapons by lysogens against susceptible competitors (Bossi et al., 2003, Burns et al., 2015, Brown et al., 2006, Sousa and Rocha, 2019). Consequently, lysogens are predicted to be better than non-lysogens at invading (or defending) ecological niches (Brown et al., 2006). Lysogens typically produce temperate phage virions at a low rate through spontaneous induction of the lytic cycle in a subpopulation, releasing phage virions into the environment (James et al., 2015, Lwoff, 1953, Nanda et al., 2015). These virions can infect nearby susceptible bacteria and replicate through the lytic cycle, killing these susceptible cells and thus enabling the lysogen to outcompete its susceptible neighbours. The utility of temperate phage weapons has been shown both in the lab and within animal infection models (Bossi et al., 2003, Burns et al., 2015). Indeed, temperate phages are thought to be an important determinant of the fitness of the Liverpool epidemic strain (LES) of *Pseudomonas aeruginosa* within cystic fibrosis patients (Winstanley et al., 2009), which is a polylysogen. Virions are actively produced in LES human lung infections and lysogens have been shown to be more competitive in experimental rat lung infections (James et al., 2015). As such, through their effects mediating bacterial competition, temperate phages contribute to controlling the dynamics of both host-associated and environmental microbial communities (Li et al., 2017).

Whilst there is good evidence that lysogens can be more competitive, less clear is how the various life-history traits of temperate phages contribute to this fitness benefit. Some work has begun to address these questions through mathematical modelling. For example, de Sousa and colleagues coupled individual based models with Random Forest analyses to explore how variability in traits such as the probability of lysis (the ratio of lytic to lysogenic infections), the phage induction rate (how rapidly new phage are produced) or the absorption rate (how rapidly free phage bind to susceptible cells) contributed to variability in the number of lysogenized resident bacteria (Sousa and Rocha, 2019). This work found that variability in key traits such as probability of lysis and burst size (how many phage virions are produced upon induction) tended to drive the greatest variability in lysogen fitness. However, whether and how much each individual life history traits altered the utility of a given temperate phage as a weapon remained unclear. Similarly, ordinary differential equation models demonstrated that the initial frequency of a lysogen is also an important factor in determining whether it can invade a resident population, with sufficient susceptible bacteria required within a population to generate exponential amplification of the phage (Brown et al., 2006). Together these mathematical results suggest both invasion conditions and phage characteristics are likely to play a critical role during lysogen invasion. Yet, despite this, we still lack any systematic study of what makes a good temperate phage weapon.

Here we combine mathematical modelling and *in vitro* experiments to comprehensively explore how different life-history traits and ecological context intersect to determine the utility of a temperate phage as a weapon. Our mathematical modelling confirms previous observations that different life history traits have very different effects on phage utility. However, we now find these impacts are highly dependent upon the frequency of the lysogen in the population. We find that certain phages may be beneficial under some circumstances yet detrimental in others, and that the “best” phage varies depending on initial population conditions. We confirm these mathematical predictions experimentally, using clinically relevant bacterial strains to confirm that Probability of Lysis plays a key role in determining phage utility, but in a manner that is strongly frequency dependent. Surprisingly though, these experiments reveal that an additional, largely overlooked environmental factor – nutrient availability – also has a critical role in modulating the impact of a phage upon lysogen dynamics. Together our results reveal the importance of the interplay between the properties of individual phages, their hosts, and their broader environments in determining the effect of temperate phages within microbial communities.

## Results

### Mathematical modelling allows us to explore determinants of phage utility

To understand how different phage life history traits and environmental factors impact the effectiveness of a temperate phage as a weapon, we adapted a previously published mathematical model of a single lysogen growing within a population of otherwise susceptible bacteria (Figure 1, Materials and Methods) (Brown et al., 2006). Here the underlying growth of each strain within the community is determined by its own intrinsic growth rate, *r*, and any interactions with self and other community members, each of which are assumed to be competitive and reciprocal. We assume that each lysogenic cell induces at a rate *I,* forming a latent infected cell which lyses at a lysis rate *p*, producing free phage virions. Free phage virions are produced at a burst size, *β* and are degraded (naturally lost) at rate *d*. Free phage virions can then be absorbed by susceptible cells at a rate, *a*. With a probability *L* these newly infected cells lyse, while with a probability 1-*L* they form new lysogens. Numerically solving this model over time enables us to simulate the dynamics of a given set of lysogenic and non-lysogenic strains over time.

**Fig 1.**
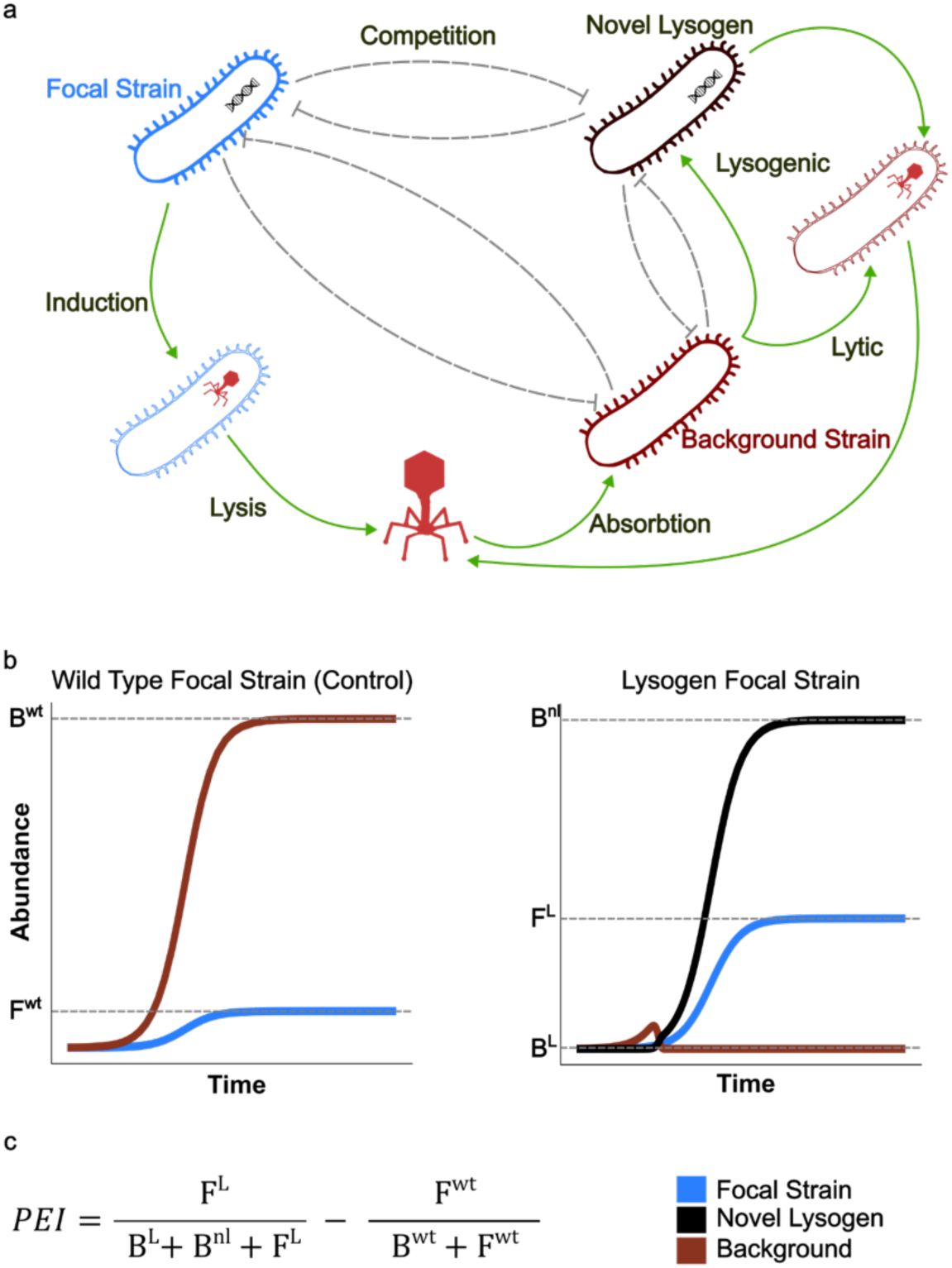
Mathematical modelling captures the dynamics of phage-mediated competition. **a**. Schematic illustrating our mathematical model. Here we focus on the dynamics of two competing strains, an initially susceptible “Background” strain (dark red), and a “Focal” strain (dark blue) which may or may not carry a temperate phage weapon. **b.** Example bacterial population dynamics when the Focal strain does (right) or does not (left) carry a temperate phage weapon. Dashed lines indicate final population abundances, which are in turn used to calculate the Phage Effectivity Index (PEI) **c.** Formula for the PEI, defined as the difference in the final frequency of the Focal strain with vs without a temperate phage.

We used this model to explore the simple scenario whereby two strains compete against one another within a niche. In our model we assumed that one strain (the *Background*, B) was always non-lysogenic, while the other strain (the *Focal*, F) could either also be non-lysogenic, or carry a temperate phage weapon. We performed these competitions under a series of different starting conditions, varying the ratio of the Focal to Background strains (F:B) to capture scenarios ranging from the initial invasion of the Focal strain into a new niche (1:99 F:B), through an evenly pitched battle (50:50 F:B), to the Focal strain defending its niche against an invading Background strain (99:1 L:B). Comparing the outcome of these competitions when the Focal strain was lysogenic to when it was non-lysogenic thus enabled us to quantify the relative advantage (or disadvantage) to a bacterium of possessing a given temperate phage, which we term the Phage Effectivity Index (PEI). We focused on this measure as it allowed us to directly capture the absolute advantage of harbouring a phage weapon (vs not) regardless of context. However, for completeness we also assessed phage effectiveness using several other metrics (SI 1-2).

### Life history traits and invasion frequency combine to shape the impact of phage weapons

Systematically varying phage life history traits in turn revealed that, as expected, each trait differed in both the direction and the magnitude of its impact on PEI. In line with previous results (Sousa and Rocha, 2019), we found that small changes in the Probability of Lysis (PoL) and the Induction Rate both led to large changes in the effect of a given phage, yet equivalent changes to traits such as the Lysis rate or Degradation Rate had virtually no impact upon PEI (Fig 2). Expanding on this previous work, our analysis now also demonstrated that increasing PoL consistently increased the effectiveness of a phage as a weapon, via reducing the rate at which novel resident lysogens were formed (Fig 2a). In contrast, increasing induction rate substantially decreased PEI – to the extent that lysogens with a high induction rate were consistently less fit than the non-lysogenic control (negative PEI, Fig 2b). That is, our analysis demonstrated that while temperate phage are beneficial on average, harbouring a lysogenic phage with a high induction rate can be worse than harbouring no phage at all.

**Fig 2.**
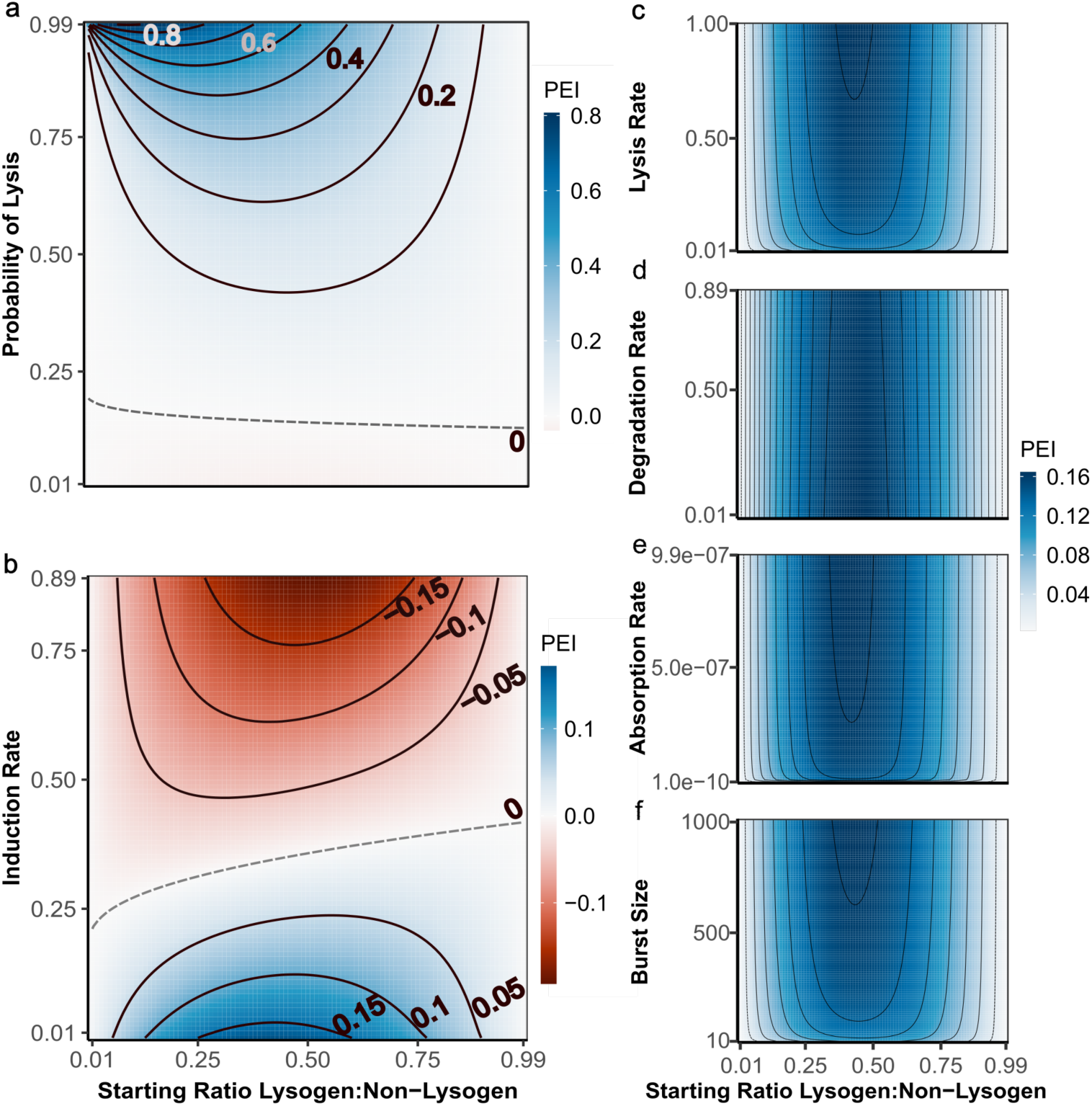
Phage life history traits and lysogen frequency jointly determine the effectiveness of a temperate phage as a bacterial weapon (PEI). Each heatmap illustrates the impact of varying a given life history trait (y-axis) and invasion frequency (x-axis) upon the effectiveness of a temperate phage weapon. Blue indicates that the phage weapon is beneficial while red indicates it is costly. Contours (black lines) show different PEI values, with dashed lines corresponding to PEI=0. Subpanels correspond to varying **a.** Probability of Lysis **b.** Induction Rate **c.** Lysis rate **d.** Degradation Rate **e.** Absorption Rate **f.** Burst size. Parameters for each simulation are given in Table 1.

**Table 1.**
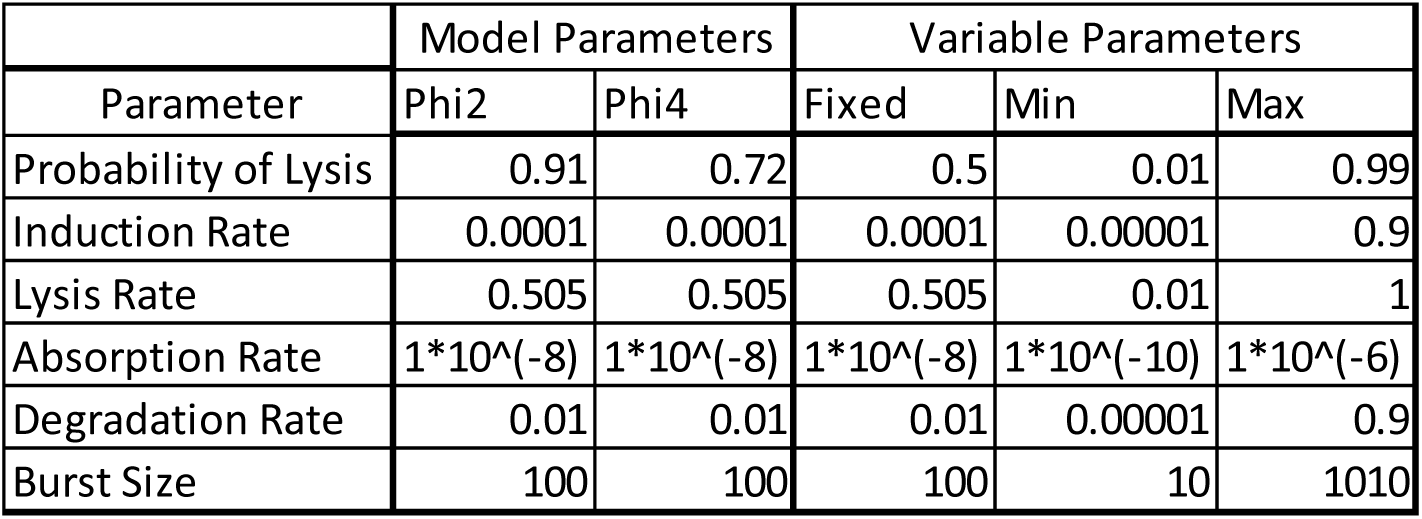
Parameters used for model simulations. “Model parameters” indicate the values used to simulate Phi2 and Phi4 for comparison against experimental data. “Variable parameters” indicate the ranges used when exploring parameter space, with the Min and Max columns showing the range of parameters explored, the Fixed column showing the values chosen when that parameter was not varied.

Our analysis also revealed that the effect of harbouring a phage on lysogen competitiveness was strongly dependent upon their initial frequency within the population. In most cases, phage provided the greatest advantage when lysogens were at intermediate frequencies. In contrast, when lysogens were initially present at either very low or very high frequencies we observed very little impact of harbouring a phage upon the success of a lysogen within a population (PEI ∼= 0). Notably however, the exact shape of this frequency dependence was in turn strongly modulated by individual phage life history traits. Phage with a high PoL were most effective when lysogens were invading from rare, while phage with a lower PoL provided the greatest advantage when lysogens were introduced to the community at higher abundances (Fig 2a). These non-linear dynamics led to situations whereby a given phage could either be costly or beneficial, depending on the starting frequency of the lysogen within the population. For example, we found that a phage with an induction rate of 0.3 could be costly to a lysogen when its frequency within the population was low, yet beneficial when lysogen frequencies were high (Fig 2b). Importantly, we observed this same phenomenon (namely life history traits and frequency jointly modulating PEI) regardless of our measure of phage effectiveness, although the exact shape of the frequency dependence varied depending upon the metric used (Figs S1, S2). Together our analyses suggest that the impact of a temperate phage weapon upon its host’s fitness is strongly dependent not only on the characteristics of the phage, but also the ecological context in which both bacterium and phage exist.

### Life history traits can have compensatory effects

Given the very different impact of different life history traits phage effectiveness, we next sought to investigate how these traits might modulate one another’s effects. More specifically, we explored whether other life history traits might be able to mitigate the detrimental effect of phages with high induction rates. Regardless of starting frequency, almost all traits were able to mitigate for high induction rates, however, some were more effective than others (Figure 3, SI 3). As expected, increasing the PoL ameliorated the impact of a high induction rate, in some cases switching the PEI of individual phage from negative to positive (Fig 3a). More surprisingly, traits such as absorption rate and lysis rate, which had very little effect on PEI when varied in isolation, also had large mitigating effects on PEI (Fig 3b,c). That is, small increases in the absorption rate could switch a high induction rate phage from being costly to beneficial. The only trait, which did not have a strong mitigating effect, was degradation rate (Fig 3d). Together, the traits able to mitigate for high induction rates appeared to be those which increase the killing of competitors, either directly (via increasing the rate or probability of lysis) or indirectly, via increasing the rate at which novel lysogens can be formed. In other words, our analysis suggested that lysogens can bear the burden of a high induction rate phage, provided this high induction also enables them to rapidly infect and kill competitors and thus clear space within their niche.

**Fig 3.**
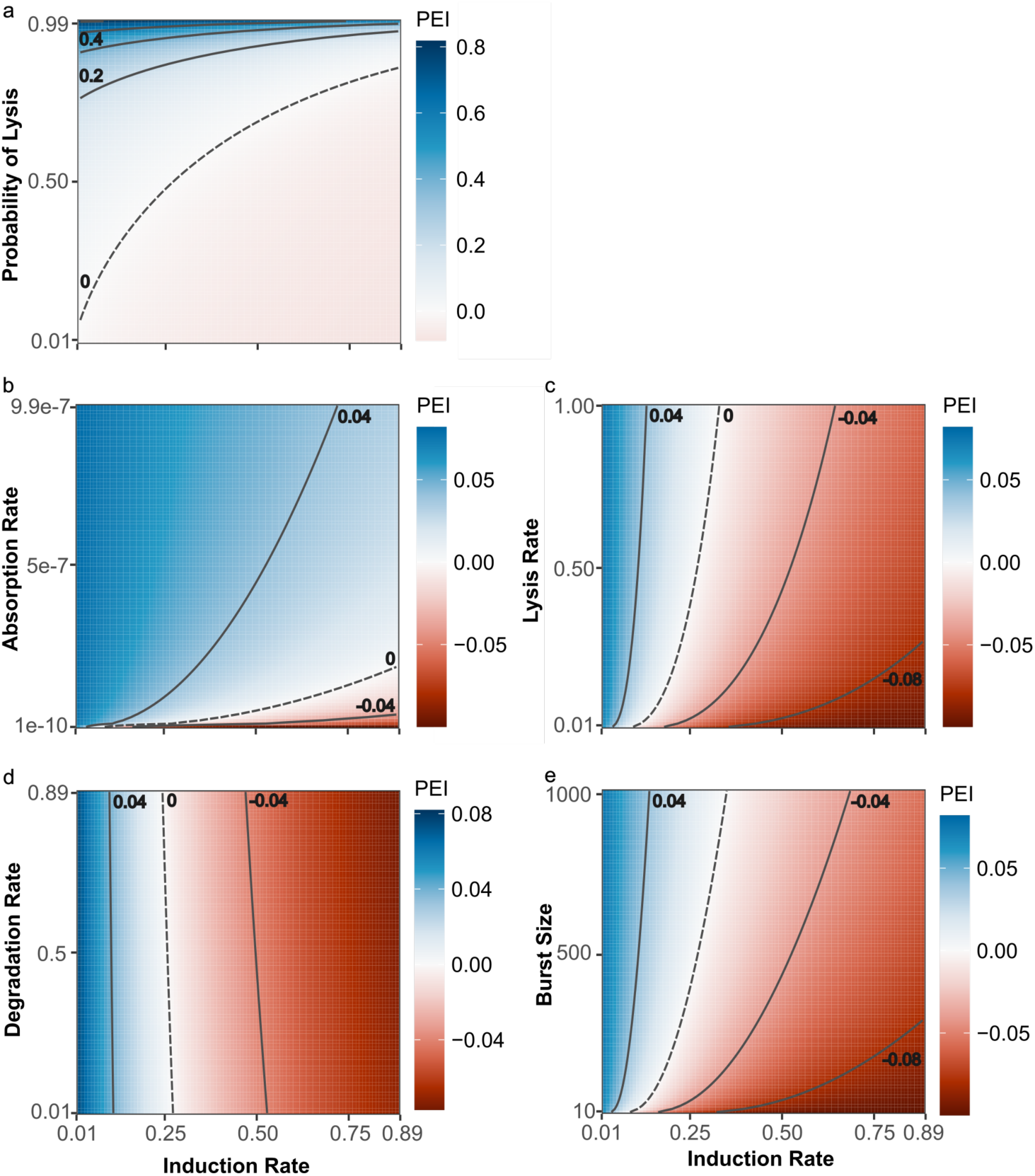
Interplay between different life history traits allows some traits to be compensatory. Here each heatmap illustrates how varying different life history traits (y-axis) against different induction rates (x-axis) changes the efficacy of different phages as weapons. Blue shows that the weapon is highly effective whereas red indicates the weapon is costly. Contours (black lines) show different PEI values, with dashed lines corresponding to PEI=0. Subpanels correspond to **a.** Probability of lysis, **b.** Absorption rate, **c.** Lysis rate, **d.** Degradation rate, **e.** Burst size. All simulations are performed with a Focal:Background starting ratio 0.1:0.9, all other parameters are given in Table 1.

### The “optimal” phage varies depending on initial population conditions

Our mathematical model suggested that the overall effectiveness of a temperate phage as a weapon is determined by a complex interplay between different phage life history traits and the initial frequency of its corresponding lysogen within a population. This led us to wonder whether this interplay between different parameters could lead to scenarios whereby the “optimal” phage weapon varied depending upon the ecological context in which a given competition was occurring. To investigate this, we randomly generated 50 different phages, each with a different unique combination of life history traits, then competed the corresponding lysogen of each phage against a non-lysogenic partner at a series of different starting frequencies. At each frequency we ranked the phage by their PEI, with 1 representing the “most useful” phage (highest PEI), and 50 representing the “least useful” (lowest PEI), such that changes in phage rank with frequency would indicate scenarios where the “optimal” phage weapon changed under different conditions.

We observed multiple PEI rank changes across starting frequencies (Fig 4a). Some phages steadily dropped in rank with increasing starting frequency – suggesting that these phages conferred a particular advantage over others when rare, but not when common. Other phages showed the opposite pattern, slowly rising through the ranks as lysogen starting frequency increased, while for some phages their PEI rank did not vary with starting frequency. To identify whether there were trends in these PEI rank changes, we repeated our ranking analysis 100 times, creating 100 sets of 100 random phages. We then quantified the number of phages that changed in PEI rank at each starting frequency for each phage set, allowing us to determine a measure of average Rank Volatility. Here a Rank Volatility ∼= 0 means that the relative utility of different phages does not vary with changes in lysogen starting frequency, whereas a Rank Volatility >> 0 implies small changes in starting frequency lead to large scale reordering of the relative utility of different phages as weapons. Our analysis revealed a clear inverse correlation between starting frequency and PEI rank volatility (fitted linear regression model Rank Volatility = 62.46 – 51.65*(Starting Ratio), t(998)=-59.25, p < 0.001, R^2^ = 0.78). Small changes at low starting frequencies (e.g. from 0.01 to 0.02) were highly likely to alter the relative utility of different phage weapons. Conversely, at high starting frequencies the relative utility of different phages remained relatively stable. Altogether, our analysis suggests that indeed different phage weapons are better in different environments, with subtle differences in individual phage life history traits having substantial impacts on whether and how a lysogen initially invades a population, but mattering less once lysogens have already established.

**Fig 4.**
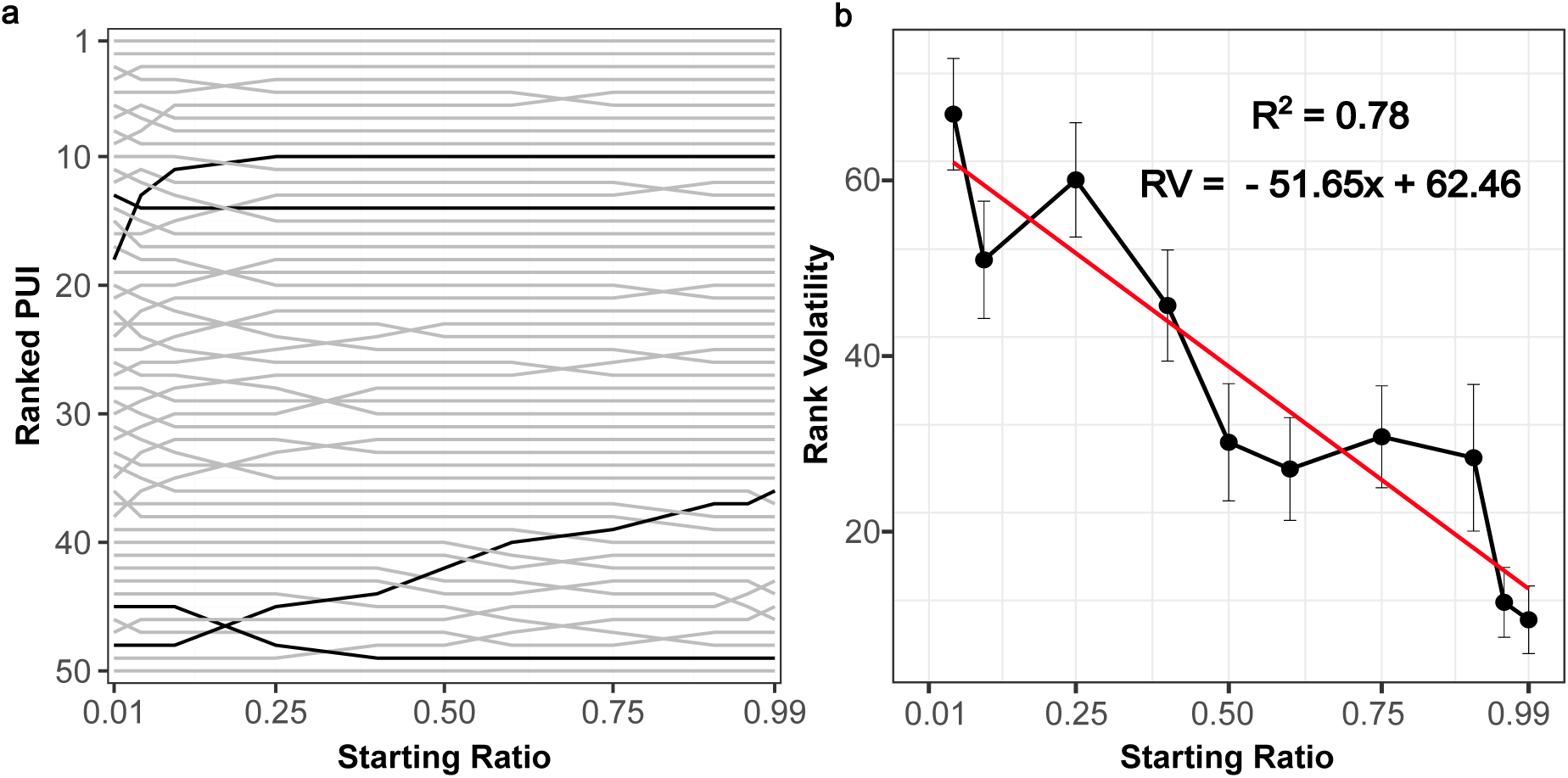
The utility of different phages changes at different starting ratios. **a**. Graph of the ranked utility (y-axis) of 50 randomly generated phages at different starting ratios (x-axis). Rank 1 corresponds to the most effective phage weapon, rank 50 to the least effective. Crossing of lines indicates that the relative effectiveness of different phage weapons is different at different starting ratios. **b.** Graph of the average number of rank changes (rank volatility – y-axis) across starting ratio (x-axis) averaged from 100 repetitions of Panel A, red line shows the fitted linear regression model, indicating a negative correlation between the rank volatility and the starting ratio. Parameters for each simulation are given in Table 1.

### In vitro experiments confirm the predicted impact of PoL and invasion frequency on phage weapon effectiveness

Our mathematical modelling enabled us to predict PEI as a function of life history traits, and initial lysogen frequency. We next sought to test the accuracy of these predictions using a simple in vitro experimental system, competing strains of *Pseudomonas aeruginosa* PAO1 with or without each of two different temperate phages (LESB58<12 or LESB58<14). First, we quantified key life-history parameters for these phages and their corresponding lysogens, revealing that the lysogens only significantly differ in their probability of lysis, with Phi2 having a higher PoL than Phi4 (Fig 3A-E). We then used these life-history data to qualitatively parameterise our model (Materials and Methods).

Our model predicted that Phi2 should consistently display a significantly higher PEI than Phi4, but that the difference in the effectiveness of each phage compared with both a susceptible strain and one another should be greatest at intermediate starting frequencies. Next, we tested these predictions experimentally by competing the lysogens against non-lysogenic PAO1 across a range of starting frequencies. Our experiments showed a good qualitative and quantitative match for each phage (Fig 3F), with Phi2 showing higher PEI than Phi4, and the PEI for both phages peaking at intermediate starting frequencies (0.1 for Phi2 versus 0.5 for Phi4). These results confirmed our specific predictions regarding the impact of PoL and initial conditions on PEI, and more generally suggested our simple model could accurately predict phage utility.

### Microbial dynamics suggest nutrient availability also shapes lysogen competition dynamics

While our experimental data qualitatively matched our mathematical predictions, we observed an interesting discrepancy between the underlying population dynamics predicted by our models and those we observed experimentally. Specifically, in control experiments when the GFP labelled non-lysogen PAO1 focal strain was introduced to a dTomato labelled PAO1 resident population, the overall bacterial density dynamics (measured by OD600 absorbance) followed near identical classic logistic growth dynamics regardless of the focal strain’s starting frequency (Fig 6a). In contrast, when the focal strain was a lysogen, the density dynamics of the overall population varied depending upon the focal strain’s starting frequency compared to a non-lysogenic control (Fig 6b,c, OD ∼ Strain x Ratio, F_5,48_ = 70.77, p<0.001). As the focal lysogen’s starting frequency reduced, populations achieved lower final densities (Figs 6d-f), and also showed stronger initial reductions in density early in the growth curve. In our simulations, however, we observed no effect of lysogen starting frequency on final population density (Fig 6g), suggesting that our model (and thus previous models) were missing a critical component of the ecology of phage-mediated microbial dynamics.

We hypothesised that this impact on final densities could be driven by the “seasonal” batch culture nature of our experiments, whereby each population is growing in an environment with a fixed starting level of nutrients that will be steadily consumed over time. In contrast, our mathematical model is based upon the classic generalized Lotka Volterra equations (Lotka, 1920, Volterra, 1926), which are known to be best suited to continuous culture conditions such as those of chemostat, but are less well-suited to batch culture environments (Picot et al., 2023). To explore whether this might be the case we altered our initial model such that the growth of each strain was now dependent upon the explicit uptake and utilization of a single nutrient within the medium (Fig 6h). Importantly, our goal here was not to capture any one specific metabolite within the environment, as we could not easily determine precisely which individual (or multiple) metabolites were being utilised within our system. Rather, we aimed to qualitatively capture one potential mechanism that might be controlling overall population dynamics.

Our nutrient-explicit model could qualitatively reproduce each of the characteristic density dynamics observed within our experiments – reproducing the correlation between starting frequency and final density, and the “dip” in densities observed at approximately 8 hours (Fig 6e). Crucially, this alteration to our model did not alter our predictions regarding phage effectiveness, producing near-identical relationships between life history traits, lysogen starting frequency, and PEI (SI 5). While our new model only captures one potential scenario, these results lead us to suggest that when lysogens are initially rare, susceptible cells are able to grow for some time – consuming nutrients within the environment – before they are infected and ultimately killed by the phage. These susceptible cells may therefore act as a nutrient sink, reducing the energy available to both the invading lysogen and newly lysogenised surviving cells, and thus limiting the final density the total population can reach. In contrast, when lysogens dominate the population, and/or susceptible residents are not killed by phage infection (as in the case of the control competitions against a non-lysogen strain) there is less potential for phage-mediated killing to remove consumed nutrients from the environment, enabling higher final cell densities. Further experimental evidence will be required to fully assess the validity of our new model. Nonetheless, our results further support our conclusion that ecological context (both in terms of starting frequencies, and factors such as nutrient availability) can play a critical role in determining the utility of a temperate phage weapon and its subsequent impact on overall community dynamics.

## Discussion

By some estimates most bacteria are lysogens (Touchon et al., 2016). Although prophage regions can degrade through evolution (Bobay et al., 2014), deactivating the phage and preventing lysis, many bacterial genomes retain active prophage(s) for long periods of time despite the risk of imminent death (James et al., 2015). A case in point is the LES, which even after decades in the lung of CF patients maintains prophage capable of inducing lysis and releasing phage virions (James et al., 2015). The only plausible explanation for the long-term maintenance of lysogeny by bacteria is that the temperate phages they habour provide them with a fitness benefit (Davies et al., 2016, Feiner et al., 2015). One potential benefit is that the temperate phages act as self-amplifying weapons that give lysogens a competitive edge by killing susceptible competitors. As such, temperate phages boost lysogen fitness both during the initial colonisation and the subsequent defence of ecological niches, with evidence for this effect accumulating from a diverse range of systems and environments (Li et al., 2017, Brown et al., 2006, James et al., 2015, Burns et al., 2015). Yet despite their ecological importance, precisely what makes an optimal temperate phage weapon has remained unclear. Here we combined simple mathematical models and experiments to disentangle how both phage life history traits and ecological context shape the effectiveness of temperate phage as biological weapons.

Consistent with previous work, our results suggest the optimal phage weapons are typically those with a low induction rate but a high probability of lysing competitor cells (Sousa and Rocha, 2019). That is, the best weapons are those that kill the lysogen as little as possible while rapidly killing competitors and minimizing the formation of new lysogens resistant to the weapon. However, our results also reveal that phage effectiveness is a nuanced and multi-dimensional property. Detrimental phage weapons (such as those with high induction rates) may be rendered beneficial through small changes in other life history traits (such as absorption rate), creating the potential for intricate evolutionary trajectories. More fundamentally, we find that the effectiveness of a phage weapon is not universal, but rather is contingent upon ecological context. The same phage weapon may be beneficial or costly depending upon the starting frequency of its lysogen within a population, and the relative ranking of different phage weapons switches with different starting conditions. While we have yet to directly compete different lysogens against one another, our work suggests these context dependent determinants of a phage weapon’s effectiveness may be an important contributor to the large diversity in temperate phage life history traits observed in nature. Another facet to this is while low induction rate may be beneficial for the bacteria, it may not be for the phage as it limits spread and abundance. How this conflict of interest plays out over long evolutionary periods is again another grey area.

An unexpected finding was the key role of nutrients in modulating the interplay between phage life history traits, initial conditions, and overall population dynamics. Comparing our simulations and experiments suggests that, in the presence of lysogens, the consumption of nutrients by non-lysogenic cells that are subsequently lysed effectively removes energy from the system, and thus generates a negative relationship between initial lysogen frequency and final population densities. While this reduction in population density did not appear to affect the overall utility of a given phage weapon in our simple two strain competitions, we suspect these dynamics could become much more important in broader context. For example, most microbes inhabit multispecies communities, competing and cooperating with numerous other taxa; as such, if lysogen invasion reduces the abundance of one taxon this disturbance may have a knock on effect upon many other taxa (Coyte et al., 2015). Understanding how these phage-nutrient dynamics play out within a broader community or meta-community model is therefore an important future avenue for research.

To explore systematically the determinants of phage effectiveness here we have used relatively simple ecological models. The advantage of these simple models is that we can screen vast numbers of temperate phage life histories in high-throughput *in-silico* experiments. As a consequence, we have omitted a number of features such as density dependent phage induction, and de novo phage resistance that are known to occur within natural systems (Erez et al., 2017, Wright et al., 2019, Rendueles et al., 2023). Understanding how these added layers of complexity modulate the effectiveness of phage weapons is therefore an important open question. However, it is notable that despite their simplicity our models could accurately recapitulate our core experimental results. This suggests that our overarching predictions – namely that the effectiveness of a given phage weapon depends not only upon its individual life history traits but also the ecological context in which its host resides – are likely to hold under a wide range of different conditions. More generally, our results underscore the power of simple mathematical models to understand drivers of microbial dynamics.

## Methods

### Initial model analysis

We initially explored bacterial and phage dynamics using a simple mathematical model first developed by Brown *et al* (2006) (note here we rewrite the equations to take the form of the generalized Lotka Volterra equations, so as to allow easy comparison with previous microbial community models. However, this alternative functional form is mathematically equivalent to the original Brown *et al* approach of logistic growth with a shared carrying capacity). This model captures the rate of change of lysogenic and non-lysogenic populations of two competing strains, yielding the following system of ordinary differential equations,

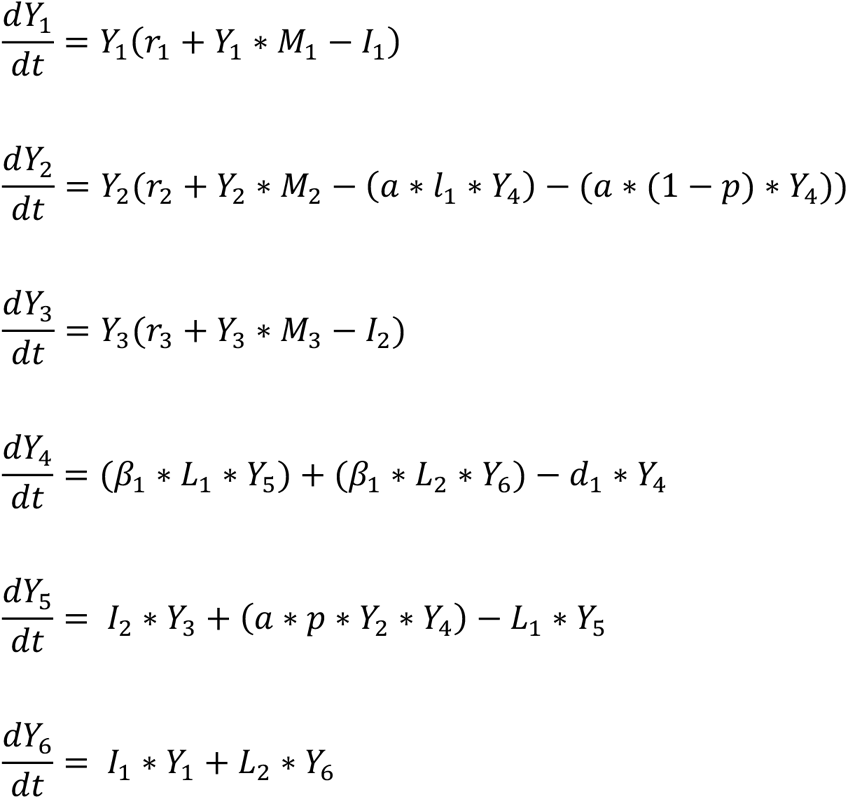

Here Y1 denotes the abundance of the Focal strain, while Y2 denotes the Background strain. Y3 is the novel lysogen which is formed when the bacteriophage (Y4) infects through the lysogenic cycle. Y5 is formed when the lysogenised Background strain is induced, or when the Background strain is infected by the bacteriophage through the lytic cycle. Y6 is the induced state of the Focal strain. Bacterial and phage life history trait parameters (*r*, *M*, *b* etc) are defined in the main text and in Table 1.

For each simulation, we initialized the system with a low-density bacterial population (initial total population size = 1*10^6^), composed of varying frequencies of Focal to Background strains (0.01:0.99). We then simulated bacterial dynamics for a fixed period of time (t = 24), which was typically sufficient for the population to stabilize, then calculated the final frequencies of the Focal and Background species in order to determine phage utility. We first explored this model with a series of systematic parameter sweeps. Specifically, we fixed a set of baseline parameters for the model (Table 1, Fixed Parameters), with each baseline parameter value chosen such that its orders of magnitude was roughly equivalent to a previous experimental measurement (Table 1). We then systematically varied each life history trait in turn across a fixed range, while holding all other parameters constant.

In our later analyses (Figs 5f, 6), we simulated dynamics using measured experimental parameters. In each case absorption rate, burst size, and degradation rate were directly measured (see below), while parameters that were challenging to measure directly such as the probability of lysis and induction rate were estimated in relative terms (see below). For example, induction rate was estimated as small and equal for both phage, as the death rate of both the lysogen and non-lysogen was similar (Fig 5a). Probability of lysis was estimated as 25% higher for Phi2 than Phi4, as experiments showed fewer colonies growing on agar containing Phi2 than Phi4. All simulations were performed using MatLab (R2022a).

**Fig 5.**
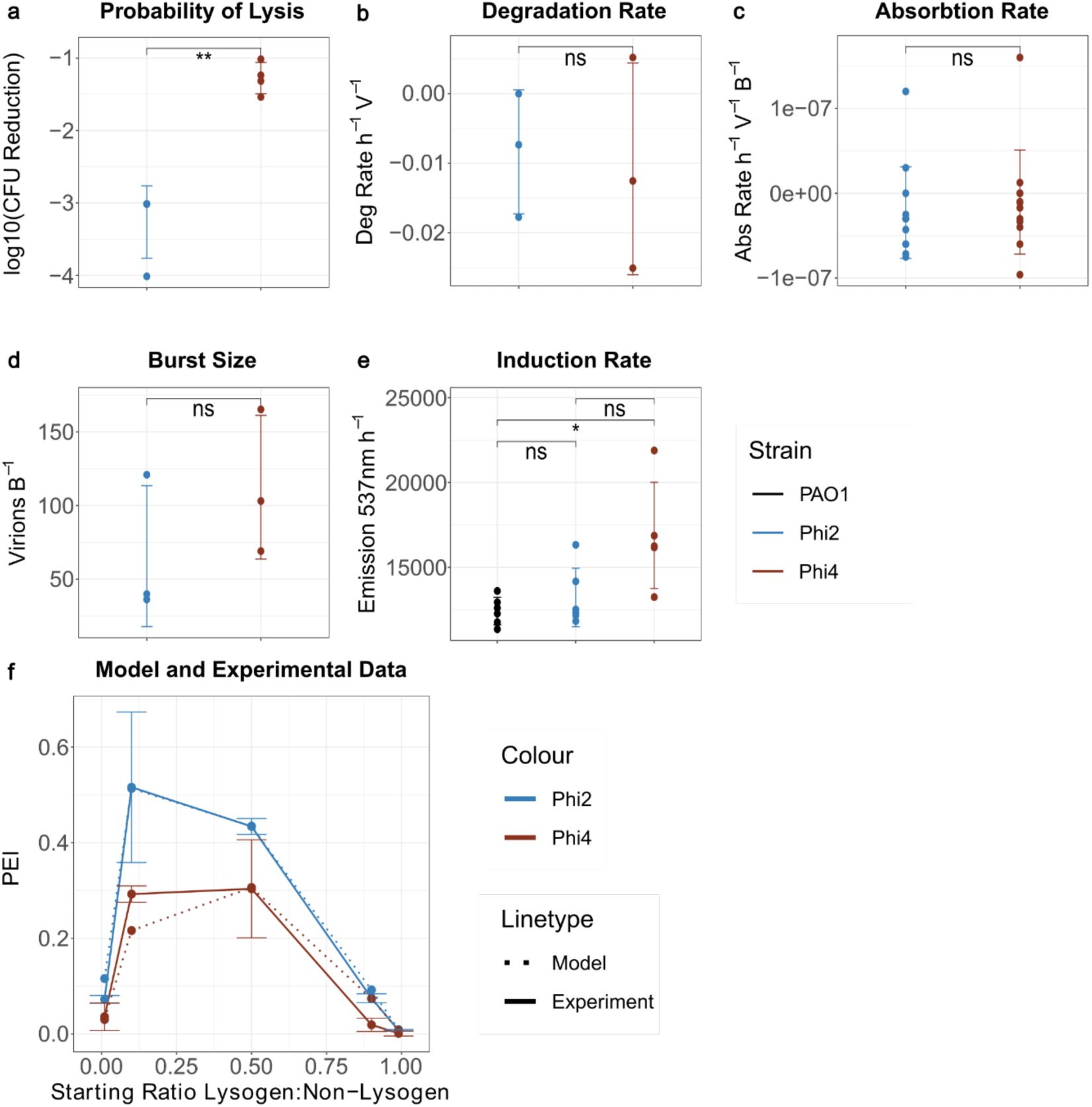
In vitro validation confirms our mathematical model can qualitatively predict the utility of different phages under different conditions. **a-e** Experimental estimates of each life history trait suggest Phi2 and Phi4 differ only in their Probability of Lysis. Subpanels correspond to **a.** Probability of lysis (p = 0.0017 **) **b.** Degradation rate (p = 0.8247 ns) **c.** Absorption rate (0.6032 ns) **d.** Burst size (p = 0.3014 ns) **e.** Induction rate (p = 0.34 ns (PAO1/Phi2), p = 0.031* (PAO1/Phi4), p = 0.058 ns (Phi2/Phi4)). **f.** Predicted (dashed lines) and measured (solid lines) Phage Effectivity Indices for Phi2 and Phi4. Predicted and measured PEIs both indicate that Phi2 is more effective as a weapon than Phi4, with this advantage most evident when lysogens are present at intermediate frequencies within the population.

**Fig 6.**
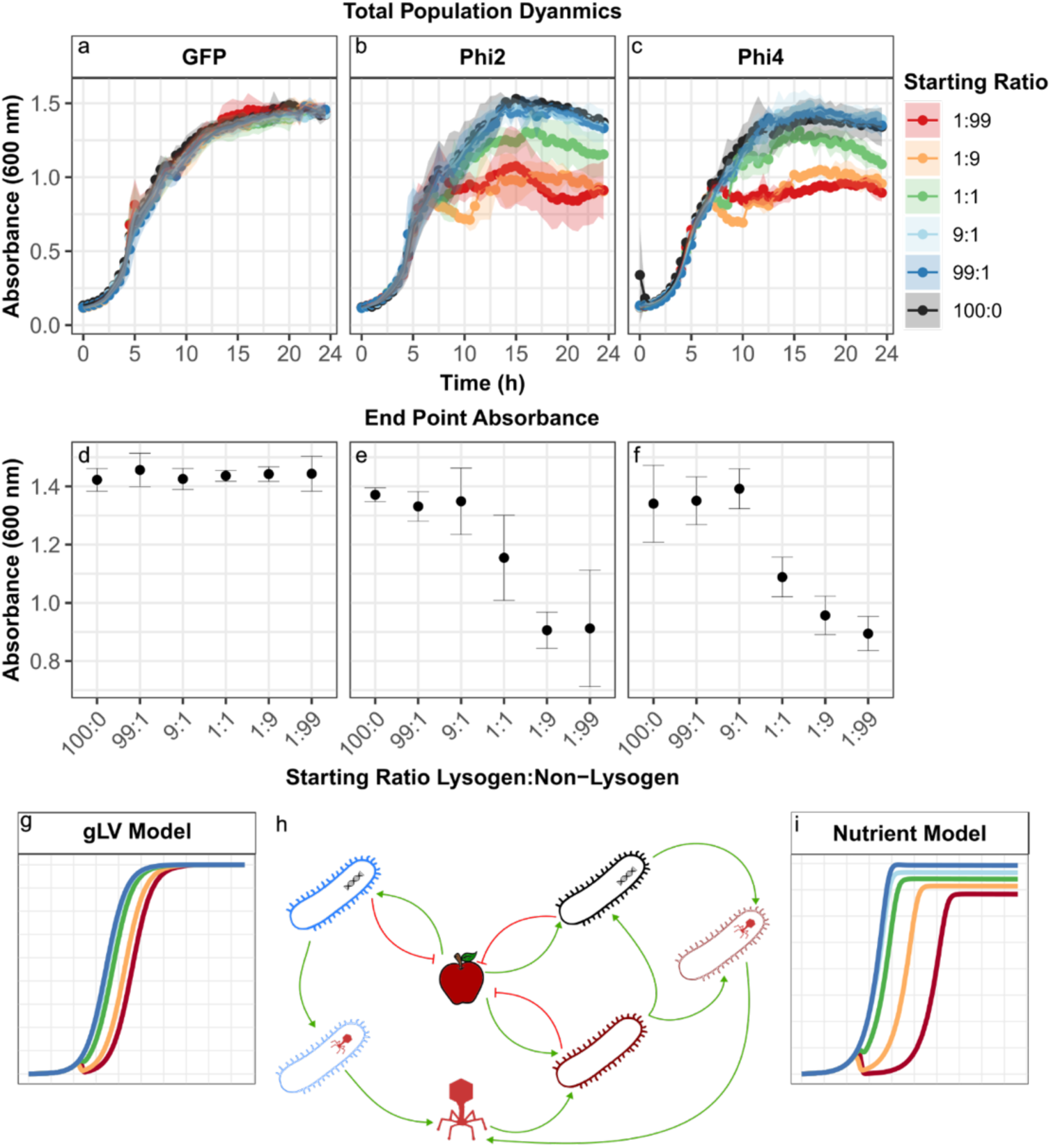
Growth curves from competition experiments reveal a relationship between initial lysogen frequency and overall population dynamics. **a-c**. Bacterial density dynamics (OD600, y-axis) for different focal strains competing against a background strain over 24-hours (x-axis). **d-f.** Comparison of the final bacterial density of the single infection control (starting ratio 100:0) to the final bacterial density of different competition experiments when the focal strain is non-lysogenic (**d**), harbours Phi2 (**e**) or harbours Phi4 (**f**). The non-lysogenic control shows no reduction in final density across the starting ratios, however lysogens exhibit a negative correlation between starting frequency and final density compared to a non-lysogenic control. **g.** Predicted growth dynamics for our original model. **h.** Schematic diagram of our model modified to explicitly include nutrient-mediated competition. **i.** Predicted population growth dynamics for the nutrient-explicit model.

### Statistics and Visualization

All graphs were plotted in R version 4.3.0 ‘Already Tomorrow’ using the ggplot package. The relationship between phage Rank Volatility (RV) and lysogen starting ratio (SR) was determined by fitting the linear model RV ∼ SR. Differences in life history traits were determined via Student’s T-test using the ggpubr package. The relationship between final population size (OD), lysogen identity (STRAIN) and lysogen starting ratio (SR) was determined by fitting the linear model OD ∼ STRAIN * SR. All statistics were computed using either base R or the ggpubr package.

### Strains, Media and Culturing Conditions

Standard cultures of lysogenised and non-lysogenised *Pseudomonas aeruginosa PAO1 (GFP,GFP-Phi2, GFP-Phi4 and dTomato),* kindly provided by C. James (Burns et al., 2015), were incubated overnight at 37°C with 180 rpm shaking in 6 ml King’s medium B (KB) in 30ml plastic microcosms. Bacterial densities were measured by plating a serial dilution of each culture onto 1.2% agar KB plates, to give colony forming units (CFU ml^-1^). Phage stocks were extracted by filtering the lysogenised bacteria overnight culture through syringe filters (0.22 µm) and stored at 5 °C. Phage densities were measured by plating serially diluted stocks onto a susceptible bacterial lawn embedded within soft agar (0.6% agar KB containing 1% GFP-PAO1).

### Absorption Rate Calculation

Absorption rate, the rate at which phages bind to bacterial cells, was calculated by mixing susceptible bacteria with phage stocks and measuring the reduction in free phage particles detected. First, 1*10^4^ PFU of each phage was mixed with 1*10^7^ CFU of GFP-PAO1 in 96-well plates. Mixtures were incubated at room temperature for 10min, then filtered using a 96-well filter plate (0.22 µm filter, centrifuged at 4500 rcf for 5 min). Phage densities were measured from starting stocks (Initial PFU) and filtrate collected after 10min incubation (Final PFU). Absorption rate was calculated using the following calculation:

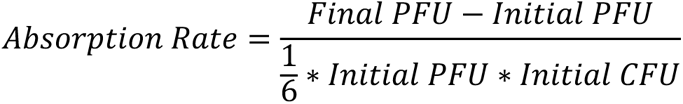

### Burst Size Calculation

Burst size, the amount of phages produced in an infection cycle, was determined by calculating the ratio of PFU:CFU at 0 and 3 hours. Phage stocks (Phi2 or Phi4) were first mixed with GFP-PAO1 at a ratio of 1000:1 bacteria:phage (1*10^7^ CFU and 1*10^4^ PFU respectively). The mixtures were grown under standard conditions for 3h after which half of each culture was syringed through a 0.22 µm filter. Phage densities were measured from starting stock (Initial PFU) and 3h filtrate (Final PFU). Bacterial densities were measured at start (Initial CFU) and from remaining 3h culture (Final CFU). The burst size was calculated using the following calculation:

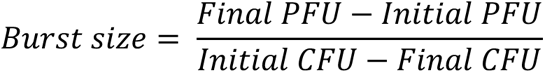

### Degradation Rate Calculation

Degradation rate (the rate phage particles become inactive) was calculated by comparing the PFU at 0 and 24 hours. Pure phage lysates were extracted by filtration (0.22 µm) from lysogensised bacterial culture grown under standard conditions. Degradation rates were calculated by incubating pure phage lysates, extracted from PAO1-Phi2 and PAO1-Phi4, without bacteria present for 24h under standard culturing conditions for lysogenised bacteria (37 °C, 180rpm), in a 1ml Eppendorf tube. The initial phage density (PFU^T0^) and endpoint phage density (PFU^T24^) were measured; the difference in PFU was transformed into a rate per hour to give degradation rate.

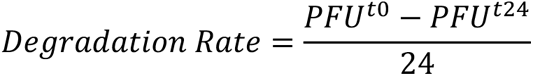

### Induction Rate

We estimated induction rate by quantifying the rate of spontaneous cell lysis. To achieve this, we quantified the production of free DNA during a bacterial growth curve using Sytox DNA stain. This method has previously been used to look at the lysis rate of cells by both bacteriocins and bacteriophages (Harhala et al., 2021, Egido et al., 2023). Here we use it as a proxy to understand if lysogens are dying significantly faster than either other lysogenised cells or non-lysogens. Individual cultures of PAO1-GFP, PAO1-Phi2 and PAO1-Phi4 (at 1×10^7^ CFU ml^-1^) were inoculated with 10µM Sytox green nucleic acid stain (Thermofisher). These were left at room temperature for 15min to enable absorption of the stain and then grown at 37°C in a plate reader (Clariostar) for 10h with shaking. The emission of Sytox was recorded (excitation 490nm, emission 537nm) at 5 and 10h post inoculation. The induction rate was estimated with the following calculation:

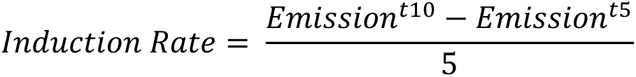

### Probability of Lysis Calculation

Probability of lysis, our estimation of the frequency of lytic vs lysogenic infections, was calculated by plating a dilution series of PA01-wt onto soft agar lawns (0.6% agar KB) embedded with each phage (1×10^8^ PFU ml^-1^) or a phage-free control (either 1% KB broth), incubated for 18h at 37C, static. Bacterial colonies were enumerated and the probability of lysis was calculated through the following calculation:

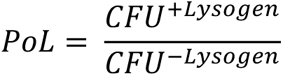

### Competition Experiments

To investigate the competitiveness of the different strains (and thereby determine phage effectiveness) we first grew our focal strains in co-culture with our background strain at a variety of different starting ratios. We then used flow cytometry to calculate the final frequency of the focal strain under each condition. Specifically, overnight cultures of lysogenised and non-lysogenised strains of bacteria were mixed at six different ratios (100:0, 1:99, 1:9, 1:1, 1:9, 1:99). Our focal strain contained a GFP fluorescent marker and our background strain contained a dTomato. Initial population ratios were validated by flow cytometry (see below). Mixtures were diluted 100-fold into KB media (6ml in 30 ml glass vial) and incubated for 24 h at 37 °C, 180rpm. After 24h the endpoint population ratios were determined by flow cytometry. To determine growth profiles, a sample of each starting mixture was also inoculated into 96-well plate with KB (1 in 100 dilution) and incubated in a plate reader (Tecan SPARK) for 24h at 37 °C, 180rpm, to track bacterial growth (OD600) every 30min.

### Flow cytometry

Chromosomal fluorescent labels (GFP or dTomato inserted within Tn7) were used to distinguish competing bacterial strains by flow cytometry and provide a population ratio. Samples were first fixed by centrifuging cultures (5000rpm, 3min) and resuspending the pellet in 4% paraformaldehyde (PFA; Thermofisher). Following 30min incubation at room temperature to complete fixation, samples were washed twice by centrifugation and resuspension in phosphate buffered saline (PBS). Diluted samples (1 in 100 in PBS) were analysed on a Cytoflex flow cytometer (Beckman Coulter), to count GFP-PA01 cells (ex. 488nm, em. 525/40nm) and dTomato-PA01 cells (ex. 488nm, em. 610/20nm). Cell counts were calculated using Kaluza Analysis (version 2.1) with electronic gates defined from single culture preparations of each strain. The population ratio was determined as the proportion of GFP-PAO1 cells within the total population (i.e., GFP-PAO1 + dTomato-PAO1).

### Nutrient-explicit model

Motivated by our experimental observations we adapted our model to explicitly capture growth on a defined nutrient source. This yielded the following system of differential equations capturing the dynamics of each bacterial population and the nutrient itself:

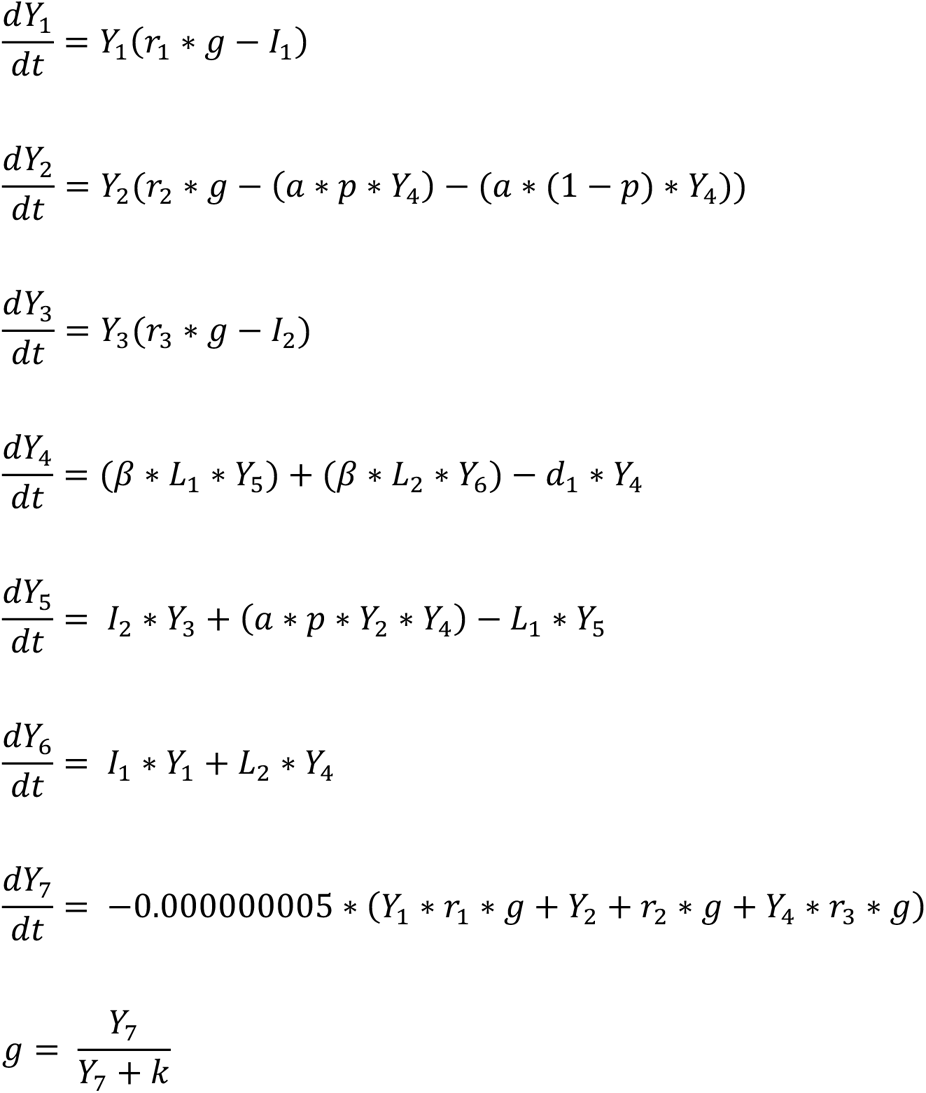

Here *Y1* to *Y6* are defined as before, *Y7* represents the amount of nutrient present within the environment, and *g* captures a saturating nutrient uptake function. In all analyses using this nutrient-explicit model phage life history traits were as before (Table 1), while nutrient uptake and conversion parameters selected to qualitatively match the growth of the non-lysogen PAO1 when in isolation. As previously, all simulations were performed using MatLab (R2022a).

## Data Availability

All code and experimental data underpinning this work can be found at https://github.com/MJNT1999/LysogenInvasionModel

## Acknowledgements

We thank Rosanna Wright for helpful comments. MJNT is supported by a BBSRC DTP Studentship, MAB by grants BB/T014342/1 and 220243/Z/20/Z, and KZC by grant 226047/Z/22/Z and a University of Manchester Presidential Fellowship.

## SUPPLEMENTARY FIGURES

**SI 1.**
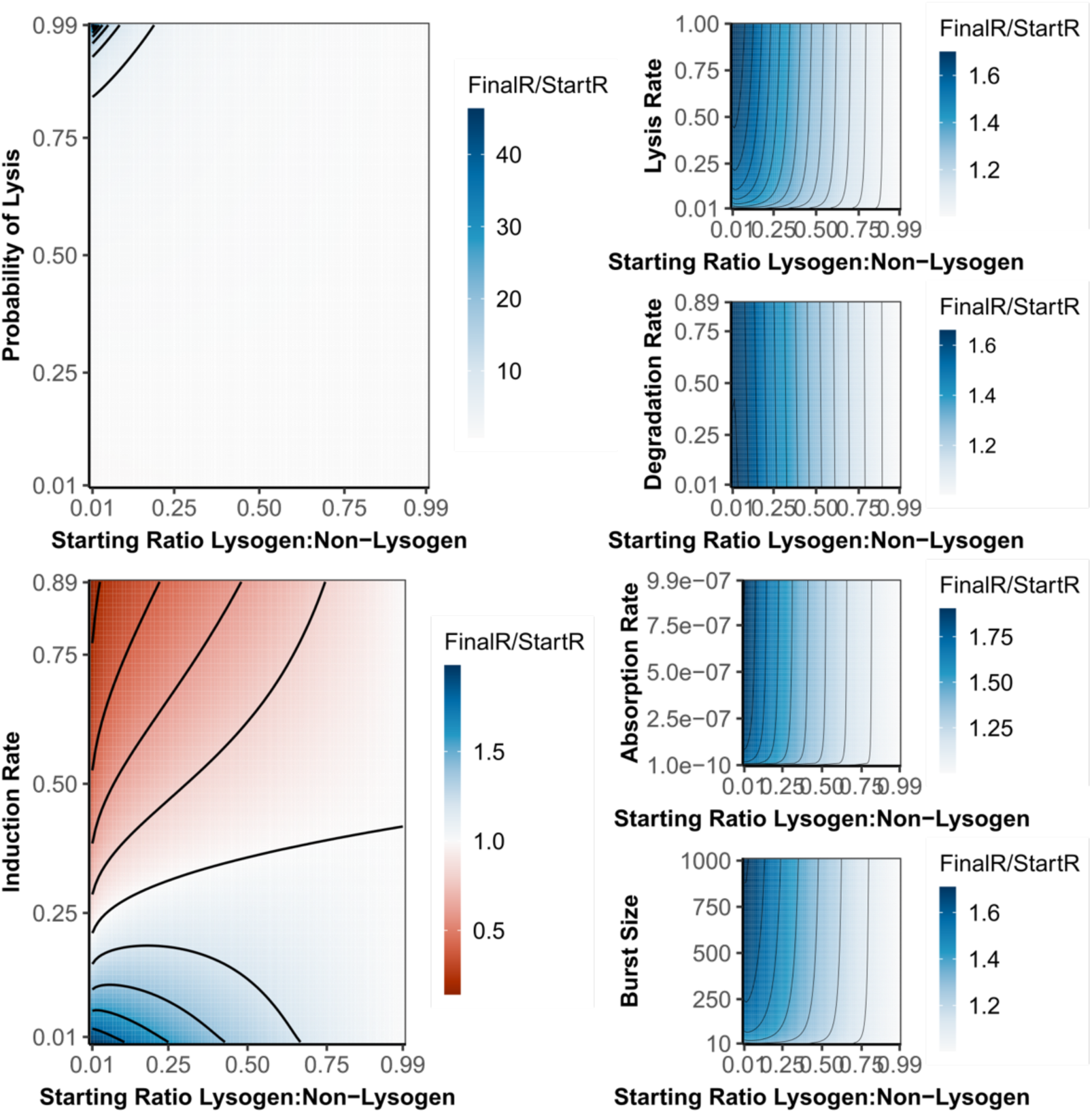
Analysing the impact of life history traits and frequency when phage effectiveness is calculated as Starting Ratio/Final Ratio. Each heatmap is the same simulations from Figure 2 but the colour now shows the PEI calculated as the final frequency of the lysogen divided by the starting frequency. As in our original analyses, we find that PoL and Induction Rate have the greatest impact on PEI, that PEI varies with lysogen starting frequency, and that the interplay between LHT and frequency leads to cases where a phage is beneficial under some conditions but detrimental under others. However, under this measure PEI is often (although not always) greatest when lysogens are rare, as this measure of competitiveness favours relatively small changes by initially rare species which in a wider community have little effect on the population.

**SI 2.**
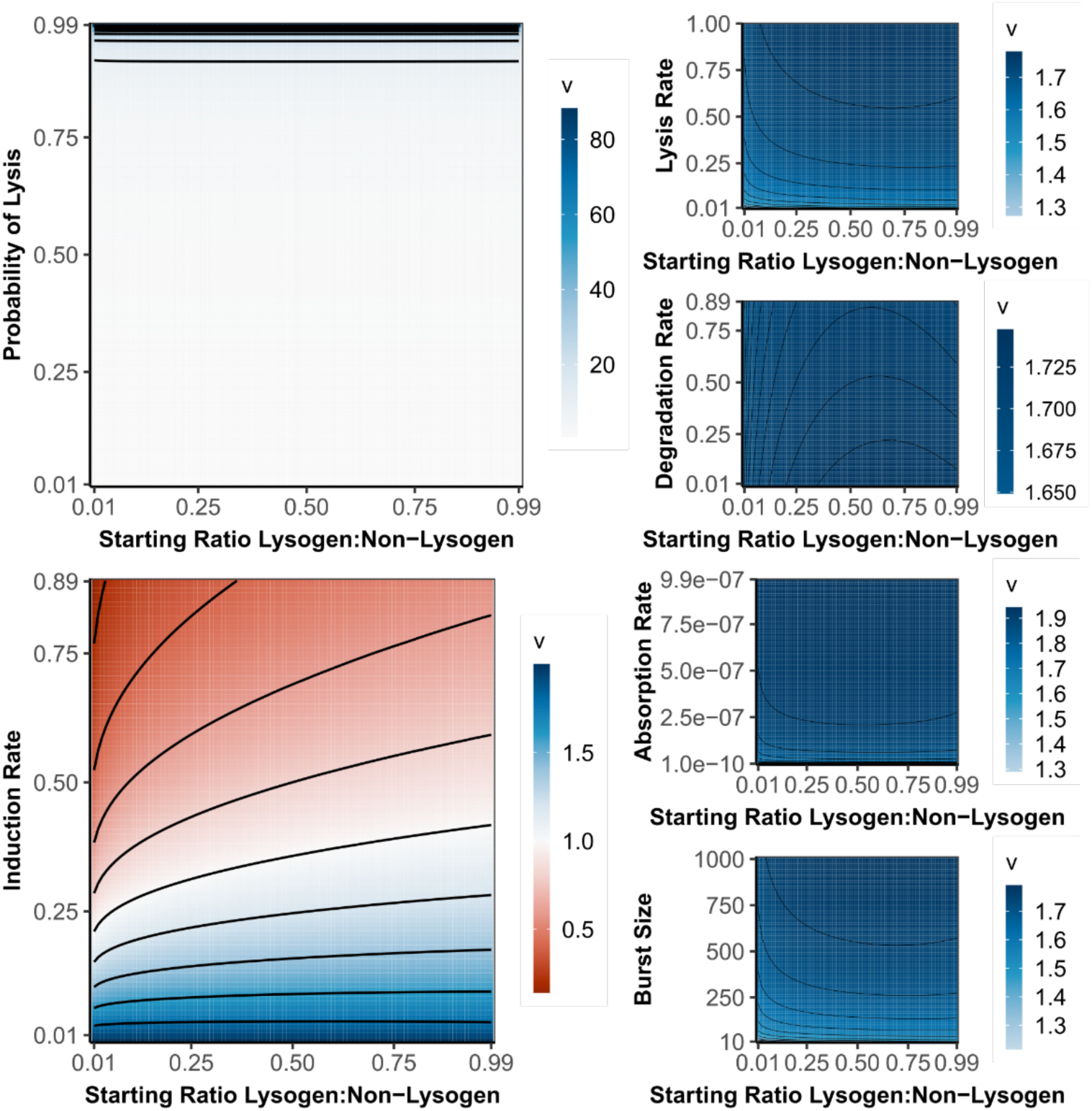
Analysing the impact of life history traits and frequency when phage effectiveness is calculated as the fitness measure, *v*. Here each heatmaps is coloured by the fitness measure v: 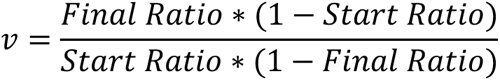 As in our original analyses, we find that PoL and Induction Rate have the greatest impact on PEI, that PEI varies with lysogen starting frequency, and that the interplay between LHT and frequency leads to cases where a phage is beneficial under some conditions but detrimental under others. However, as *v* tends to saturate quickly these effects are less prominent than with other measures.

**SI 3.**
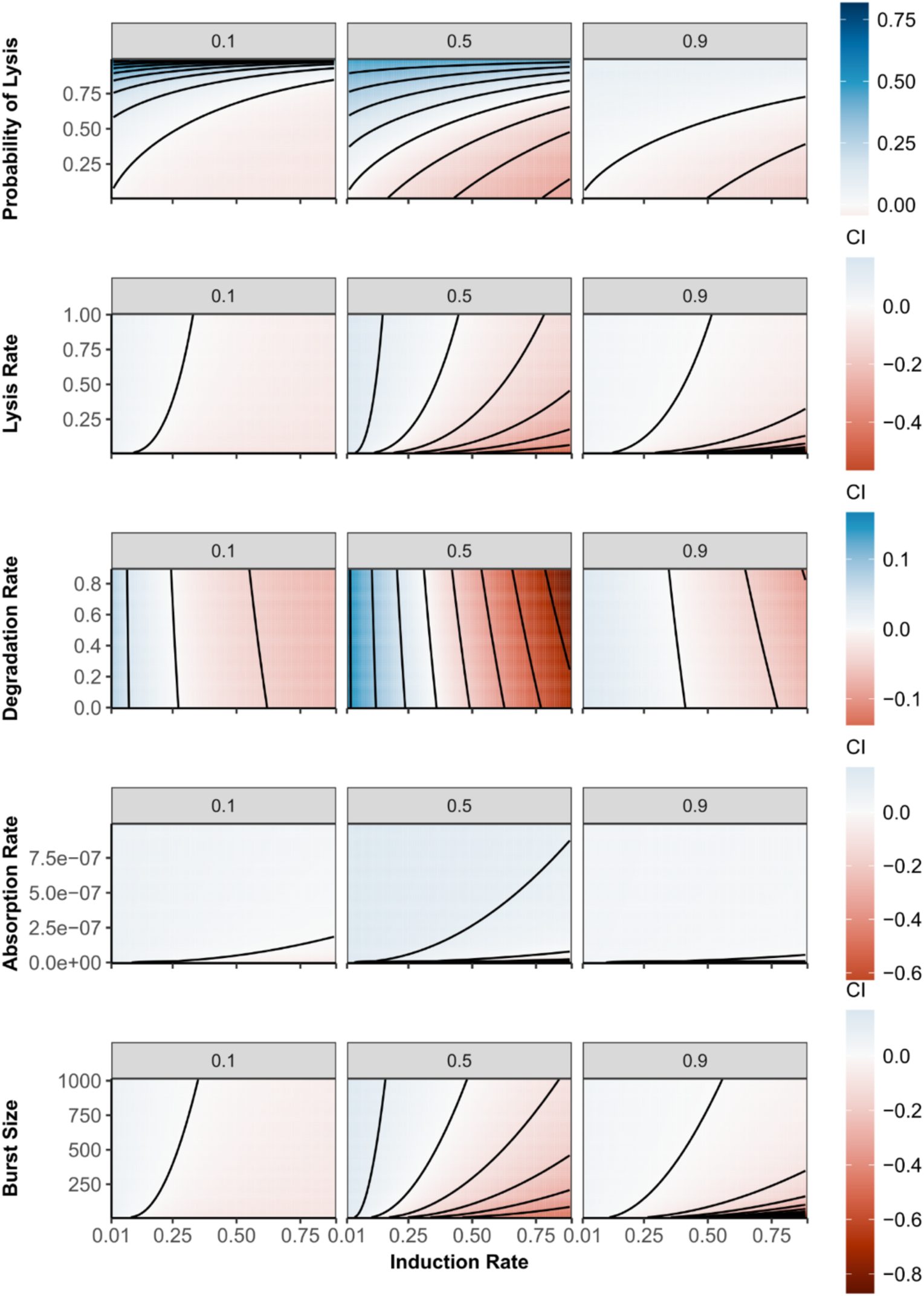
Compensatory effects of life history traits are observed at several starting frequencies. Each heatmap shows equivalent parameter sweeps to those performed in Fig.3, but with different starting frequencies of the Focal strain. While at different frequencies we see subtle changes in the dynamics, the overall relationship remains the same.

**SI 4.**
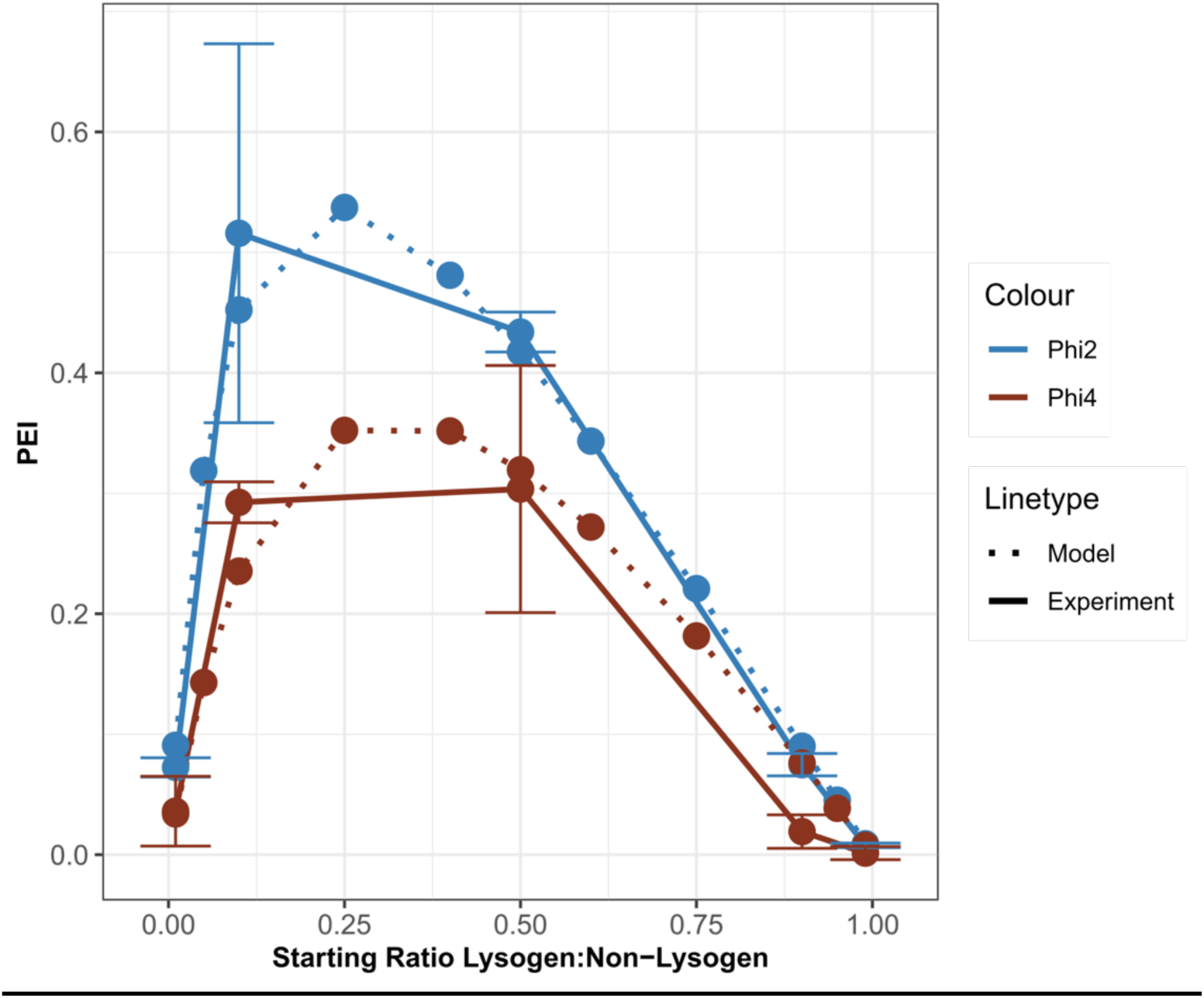
Our nutrient-explicit model can also well accurately predict our experimental results. Predicited values for the nutrient model are shown in dashed lines while experimental values are shown with solid lines. The predictions for Phi2 are shown in blue and Phi4 in red. The nutrient model also predicted Phi2 to be more competitive than Phi4 while also recapitulating the shape of the curve.

**SI 5.**
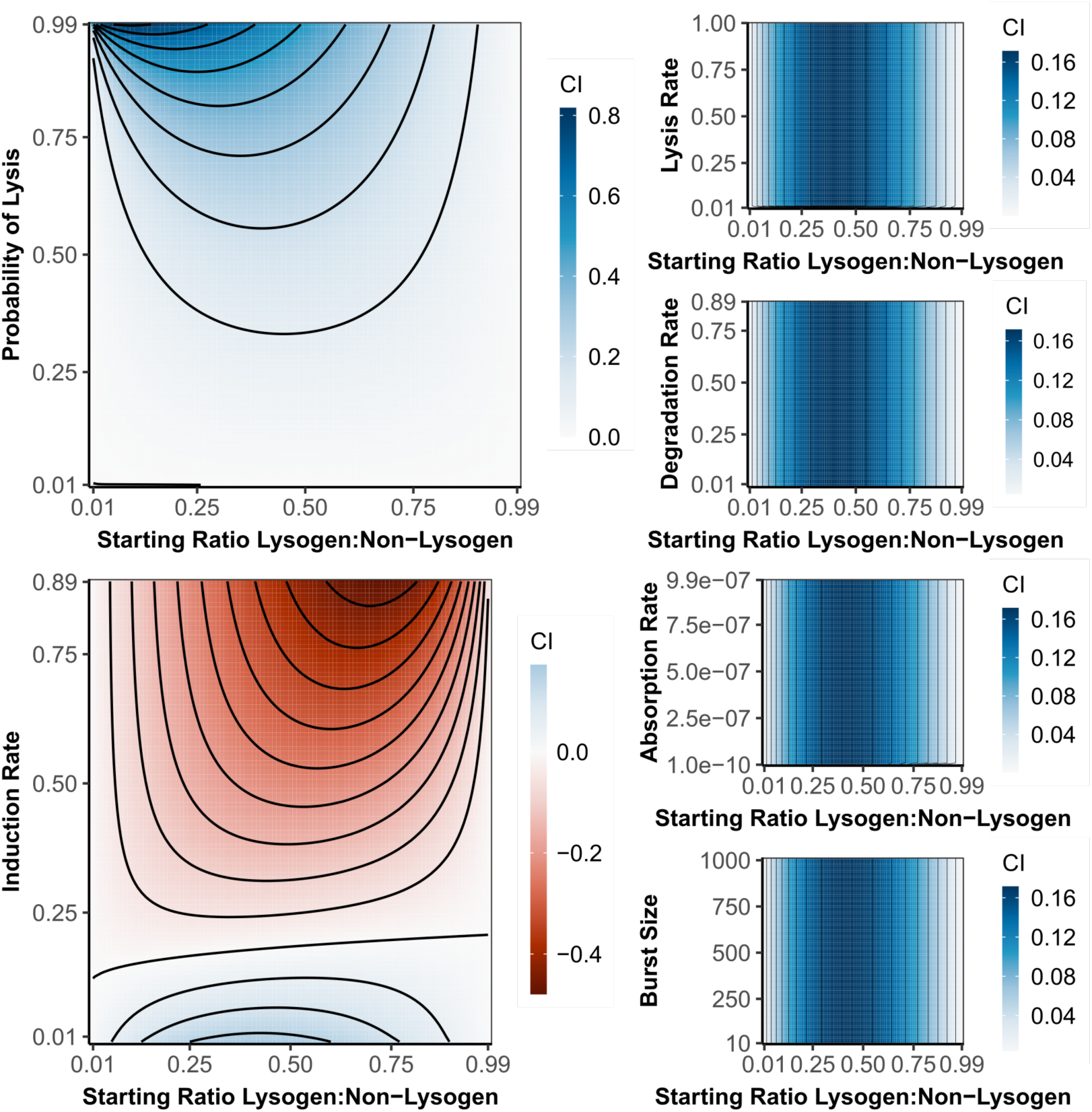
The nutrient model also recreates the relationship between different LHTs and starting frequency seen in figure 2. As before probability of lysis and induction rate have the largest effect on the competitiveness of the lysogens. While the dynamics between the models are subtly different, the overall relationships between the traits and starting frequency remains the same.

